# A Testis ER Chaperone Marks Mature Olfactory and Vomeronasal Sensory Neurons

**DOI:** 10.1101/237719

**Authors:** Ryan P Dalton

**Author notes:** Correspondence to be sent to: Ryan Dalton, Miller Institute for Basic Research, University of California Berkeley, Berkeley, CA, USA.

## Abstract

The proper folding of most secreted and membrane proteins involves interaction with endoplasmic reticulum-resident, glycan-binding chaperones. Some of these chaperones, such as Calreticulin and Calnexin, are nearly ubiquitous, while others are found only in specific cell types, presumably reflecting a role in biosynthesis of proteins specific to those cells. Herein, I have identified Calmegin (Clgn), a chaperone required for fertile spermatogenesis, as a marker of mature neurons in the olfactory system. CLGN was expressed by olfactory marker protein (OMP)-positive neurons in both the main olfactory epithelium (MOE) and the vomeronasal organ (VNO). CLGN was detected both in the perinuclear ER network and in axons. Finally expression of *Atf5*, a transcription factor required for OSN and VSN development, was both required and sufficient for robust CLGN expression in OSNs and VSNs. Together these findings establish that an ER chaperone required for sperm fertility is developmentally regulated in olfactory neurons, provide a novel marker of mature olfactory neurons, and suggest common mechanisms of secretory protein biogenesis in these cell types.

## Introduction

Mice possess two olfactory organs: the main olfactory epithelium (MOE), and the vomeronasal organ (VNO)(Dalton and Lomvardas, 2015). The MOE houses OSNs, each of which expresses a single OR allele(Buck and Axel, 1991), though there are exceptions(Liberles and Buck, 2006) (Greer et al., 2016). The VNO contains two broad classes of vomeronasal sensory neurons (VSNs) expressing different types of vomeronasal receptors (VRs). Apical VSNs express a single type I vomeronasal receptor (V1R) and basal VSNs express two type II vomeronasal receptors (V2Rs) in non-random combinations(Berghard and Buck, 1996). VRs are thought to mainly be activated by pheromones, resulting in the generation of stereotyped physiological and behavioral responses(Halpern, 1987; Dulac and Axel, 1995). The receptor gene regulation program in both OSNs and VSNs involves an initial process of receptor gene activation, followed by receptor-elicited feedback(Magklara et al., 2011; Clowney et al., 2012; Lyons et al., 2013; Markenscoff-Papadimitriou et al., 2014; Monahan et al., 2017). ORs and VRs both elicit feedback by activating the PERK branch of the unfolded protein response, a ubiquitous homeostatic signaling pathway that maintains the protein folding environment of the ER(Dalton et al., 2013). Activation of PERK drives translation of Activating Transcription Factor 5 (*Atf5*), which then promotes OSN or VSN maturation and survival, and terminates further receptor gene activation.

The discovery of these ER-centric feedback pathways was motivated by the multitude of reports indicating that chemoreceptors expressed in heterologous systems fail to productively traffick from the ER, hampering efforts to identify OR and VR ligands. A number of factors have been identified that can aid in ER exit and plasma membrane expression of ORs and VRs. Among these are receptor transporting protein 1 and 2 (Rtp1/2) (Saito et al., 2004; Bush and Hall, 2008; Sharma et al., 2017), receptor expression enhancing protein 1 (REEP1), and Hsc70t(Neuhaus, 2006), a testis-enriched member of the Hsp70 family of chaperones. These factors appear to increase cell surface expression for some, but not all, ORs. In contrast to findings on ORs, V2R trafficking appears to be negatively regulated by the ubiquitous soluble chaperone calreticulin (Calr)(Dey and Matsunami, 2011). Calr is not expressed in VSNs, having been replaced by the VNO-specific homolog Calreticulin 4 (Calr4). Depletion of Calr from heterologous cells enhances V2R surface expression. No factors regulating plasma membrane expression of V1Rs have yet been identified. Given the extraordinarily large size of the OR and VR gene families, it is likely that additional factors involved in the biogenesis of specific subsets of receptors have yet to be discovered.

It has become increasingly clear over the last several years that OR expression—as well as expression of other chemoreceptors such as taste receptors—is by no means restricted to peripheral sensory cells, being found in cell types as different as immune cells and somatosensory neurons(Li et al., 2015; Malki et al., 2015). Indeed, it was not long after the discovery of the OR gene family that OR expression was first identified in sperm(Parmentier et al., 1992). OR expression in sperm appears to be functional in at least some cases, as sperm expressing the olfactory receptor for bourgeonal, hOR17-4, as well as a number of others, exhibit calcium and chemotactic responses to their ligands(Flegel et al., 2015) (Spehr et al., 2003). Given the aforementioned requirements for functional OR expression in OSNs, it is likely that molecules specifically promoting OR biogenesis are expressed during spermatogenesis. A study of testis-enriched ER proteins therefore could aid in the identification of additional molecules involved in biogenesis of ORs and other OSN-specific proteins. Herein, I have identified Clgn, a testis-specific chaperone required for proper spermatogenesis(Tanaka et al., 1997; Ikawa et al., 1998; Tokuhiro et al., 2012), as highly-enriched in mouse olfactory organs, and I characterize in detail its expression pattern and regulation in both OSNs and VSNs.

## Materials and Methods

### Mice and Strains Used

All mice were housed in standard conditions with a 12-hour light/dark cycle and access to food and water. All mouse experiments were approved by and were in accordance with University of California IACUC guidelines. All strains were maintained on a mixed genetic background. The following mouse lines have been previously described: *Eif2αS51A/S51A*(Scheuner et al., 2001; Dalton et al., 2013), Atf5-/-(Dalton et al., 2013), *OMP-GFP*(Li et al., 2004), *tetO-Atf5(Dalton* et al., 2013), (Nguyen et al., 2007).

### Immunofluorescence

IF was performed as has been previously described(Clowney et al., 2012; Dalton et al., 2013; Lyons et al., 2013). Tissue was directly dissected into OCT or dissected into 4% PFA in PBS. 14um cryostat sections were air-dried for 10 minutes, fixed in 4% PFA in PBS for 10 minutes, washed 3x5 minutes in PBS + .1% Triton-X (PBST), blocked for 1 hour in 4% donkey serum in PBST, then incubated with primary antibodies under coverslips overnight at 4C. The following day, slides were washed 3x15 minutes in PBST and then incubated with secondary antibodies and DAPI at concentrations of 1:1000 under coverslides. Slides were then washed 3x15 minutes in PBST and mounted with vectashield for imaging. Imaging was performed on Leica 700-series laser scanning confocal microscopes. The following antibodies were used: goat anti-Clgn, 1:250 (SCBT, discontinued), rabbit anti-Adcy3, 1:300 (SCBT SC-588).

## Results

### Clgn is Expressed by Mature OSNs and VSNs

In order to identify testis-enriched ER genes that are also expressed in the olfactory organs, I searched published mRNA-seq databases from mouse OSNs taken at specific developmental timepoints(Magklara et al., 2011). This search identified *Clgn* mRNA as highly-enriched in mRNA-seq datasets derived from mature, olfactory marker-protein positive OSNs. In order to confirm this expression pattern, I performed immunostaining of olfactory epithelia as well as testis from adult animals. In support of the specificity of this antibody and consistent with previous reports, CLGN signal in testis was confined to spermatocytes and early spermatids (Figure 1A-B) (Watanabe et al., 1992; Yoshinaga et al., 1999). In coronal sections of the olfactory epithelium from *OMP-GFP* (Li et al., 2004) animals, in which mature OSNs are labeled with GFP, GFP+ cells were also CLGN+, as expected from the above-mentioned mRNA-seq studies (Figure 1C-F). CLGN signal was found both in the perinuclear cell bodies and in the axon bundles (Figure 1E, 3A), suggesting that CLGN is found in the OSN ER network. CLGN immunoreactivity was also found to overlap with OMP expression in the VNO, where OMP marks mature VSNs (Figure 2A-D). Importantly, CLGN was found in both apical and basal areas of the VNO, corresponding to both major classes of VSNs. Together, these results indicate that Clgn is a highly-enriched ER marker of mature OSNs and VSNs, suggesting that it may function in the biogenesis of proteins specific to mature olfactory neurons.

**Figure 1.**
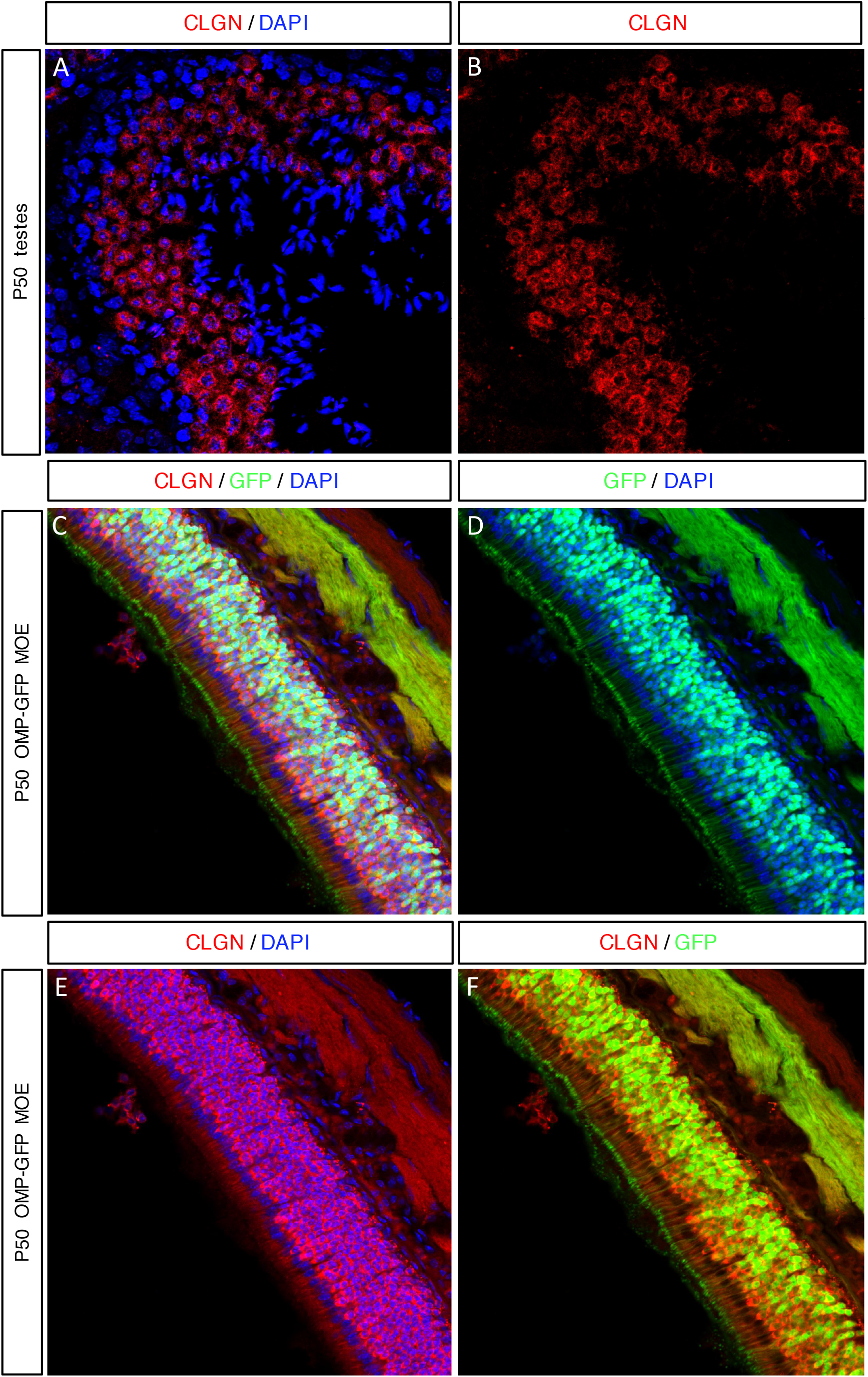
Expression pattern of CLGN in testis and olfactory epithelium. Representative laser scanning confocal microscope (LSM) images. Shown are 14um-thick sections from adult mouse testis or olfactory epithlium were used for immunofluorescence microscopy. (A-B) CLGN immunofluorescence (IF) in red and DAPI nuclear counterstain in blue reveal the expression pattern of CLGN during spermatogenesis. Heads of sperm can be seen central to the ring of CLGN. (C-F) Coronal sections from *OMP-GFP* olfactory epithelium with native GFP fluorescence in green, DAPI nuclear counterstain in blue, and CLGN IF in red.

**Figure 2.**
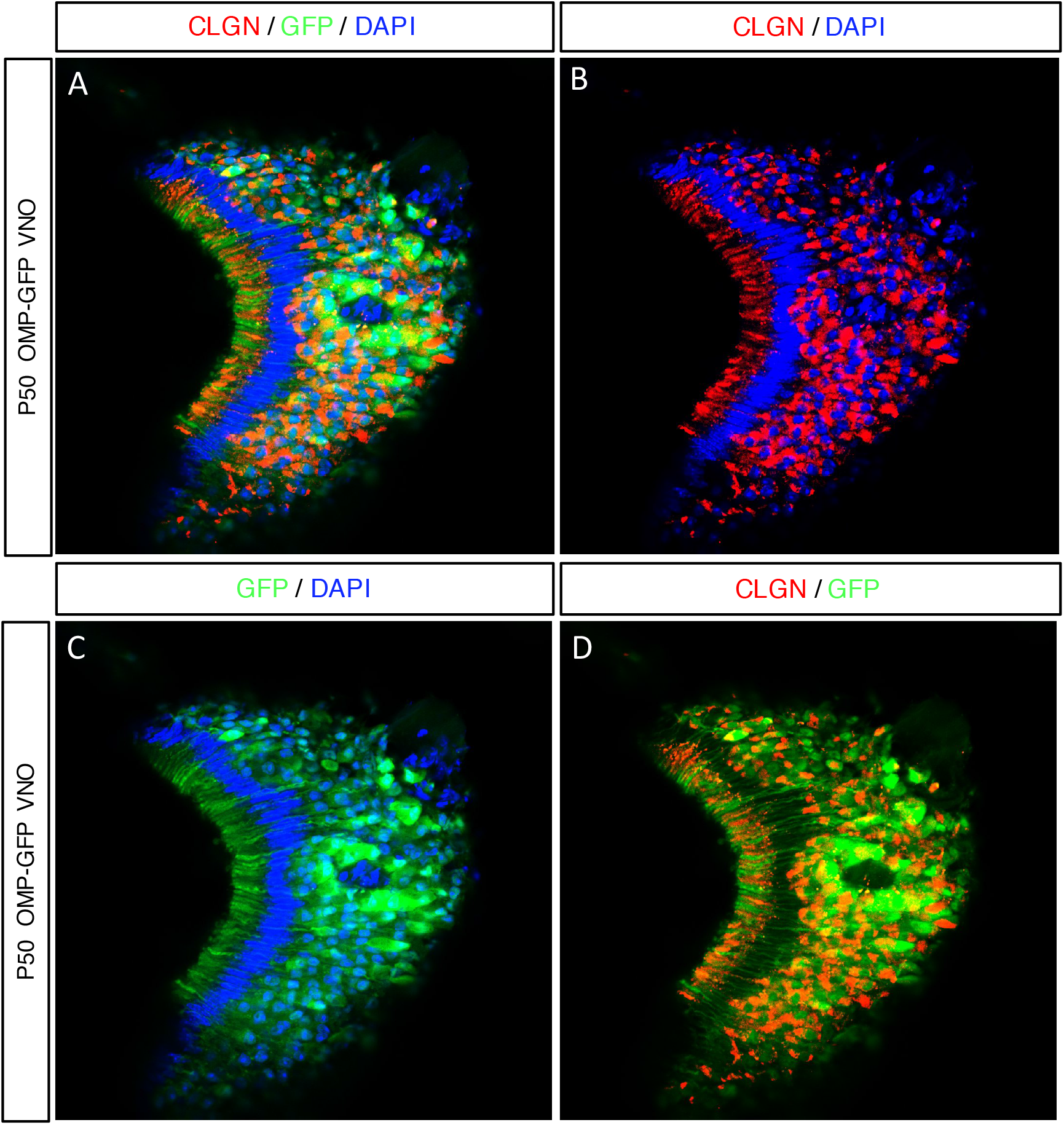
Expression patterns of CLGN in vomeronasal epithelium. Representative LSM images. Shown are 14um coronal sections from a P50 *OMP-GFP* animal with native GFP fluorescence in green, DAPI nuclear counterstain in blue, and CLGN IF in red. Apical areas are to the left.

**Figure 3.**
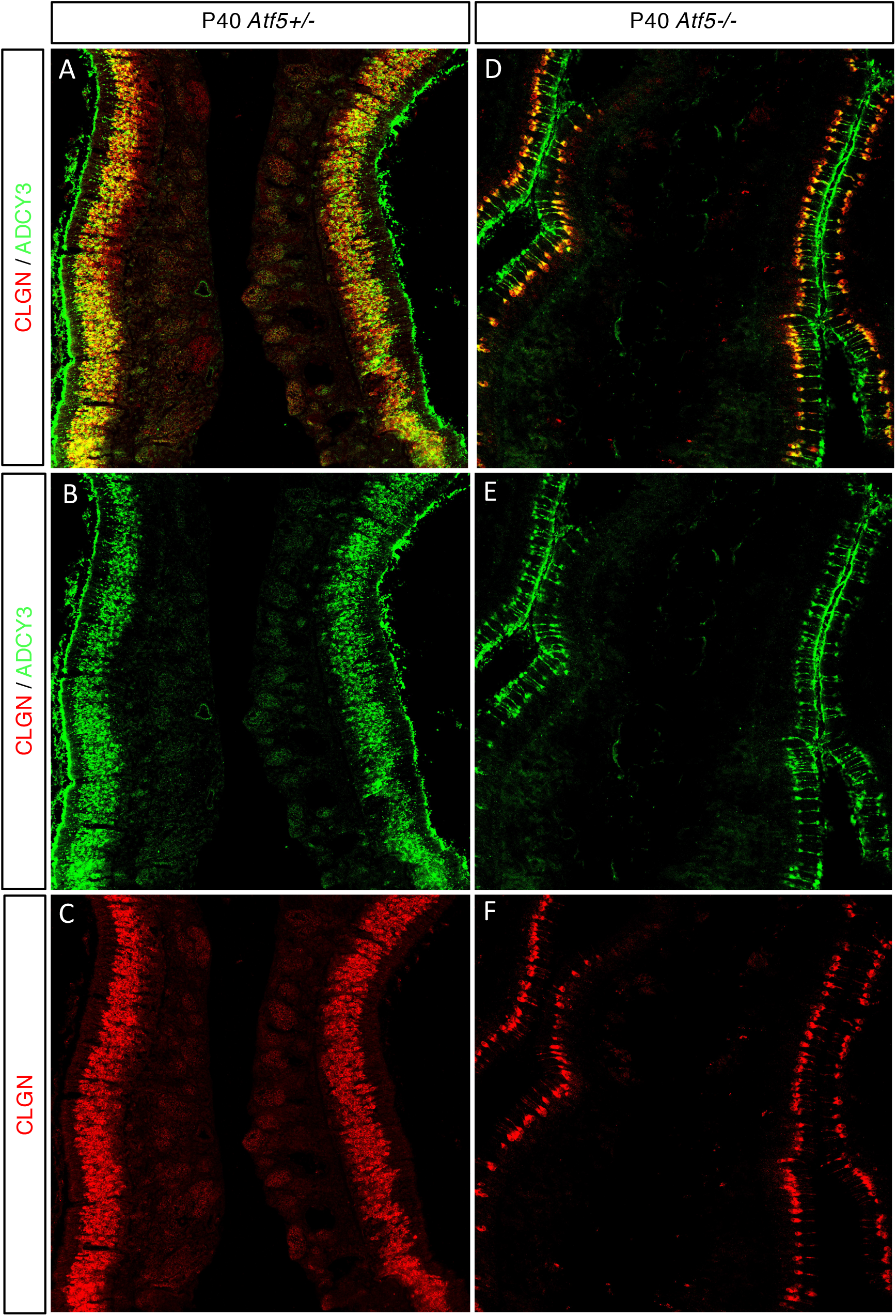
CLGN and ADCY3 are co-regulated. Representative LSM images. Shown are 14um coronal sections from P40 *Atf5+/-* and *Atf5-/-* littermates. Sections are through the septum for both genotypes to show comparison. ADCY3 is shown in green and CLGN in red. (A-C) Control animal displays an olfactory epithelium with ADCY3 and CLGN signal several cell layers deep. (D-F) The small number of persisting neurons in *Atf5-/-* are positive for both CLGN and ADCY3.

### *Atf5* is Required and Sufficient for CLGN Expression in OSNs

OR expression in OSNs is required and sufficient to drive translation of *Atf5.* ATF5 is likewise required and sufficient to promote OSN maturation and expression of the canonical OR signaling molecule adenylyl cyclase 3 (Adcy3) and the OR transporter Rtp1(Wang et al., 2012; Dalton et al., 2013). *Atf5* mutant animals display a nearly total loss of mature OSNs, with the persisting OSNs arranged in an intriguing, regular spatial pattern. In order to determine whether *Atf5* is required for CLGN expression, I first assayed ADCY3 and CLGN expression in an adult *Atf5* mutant animal. As can be seen in **Figure 3A-C**, in control animals, ADCY3 cells co-stained for CLGN, as expected from **Figure 1**. In *Atf5* mutant animals, CLGN and ADCY3 again marked the same cells, indicating that *Adcy3* and *Clgn* are co-regulated by ATF5.

Translational control of *Atf5* is exerted through activation of the PERK branch of the UPR, which drives phosphorylation of the translation initiation factor eIF2α(Watatani et al., 2008; Zhou et al., 2008). Mutation of the eIF2α phosphorylation site serine-51 to an alanine prevents *Atf5* translation in OSNs and other cell types. In order to determine the role of ATF5 in regulation of Clgn, I therefore assayed CLGN expression in eIF2α phosphomutant (*eIF2alpha S51A/S51A*) and control (*eIF2S51A/+*)(*Scheuner* et al., 2001) animals. CLGN immunostaining was performed in postnatal day 0 (P0) animals, as eIF2α mutants die shortly after birth. In control animals, in which one copy of eIF2α was intact, CLGN immunoreactivity was widespread in the apical regions of the MOE, corresponding to mature OSNs (**Figure 4A-B**). In contrast, eIF2α mutants displayed greatly decreased CLGN signal, consistent with the defects observed in *Atf5* mutants. In addition, the distribution of CLGN in mutants was less regular and more punctate (**Figure 4C-D**). In order to determine whether *Atf5* was sufficient for CLGN expression, I assayed CLGN immunoreactivity in an eIF2α mutant animal in which *Atf5* expression was rescued by a previously-published transgene(Dalton et al., 2013). As seen in **Figure 4E-F**, this rescue approach greatly enhanced the level of CLGN expression. Interestingly, the CLGN rescue appeared to be stronger in more lateral areas, suggesting that there may be zonal aspects to the control of CLGN expression. However, it is not possible with this data to rule out the possibility that the Atf5 transgene is expressed in a heterogeneous manner, resulting in rescue of CLGN expression in some OSNs and not others. This requirement and sufficiency of *Atf5* for CLGN expression supports a model in which *Clgn, Adyc3*, and *Rtp1* are co-regulated during development.

**Figure 4.**
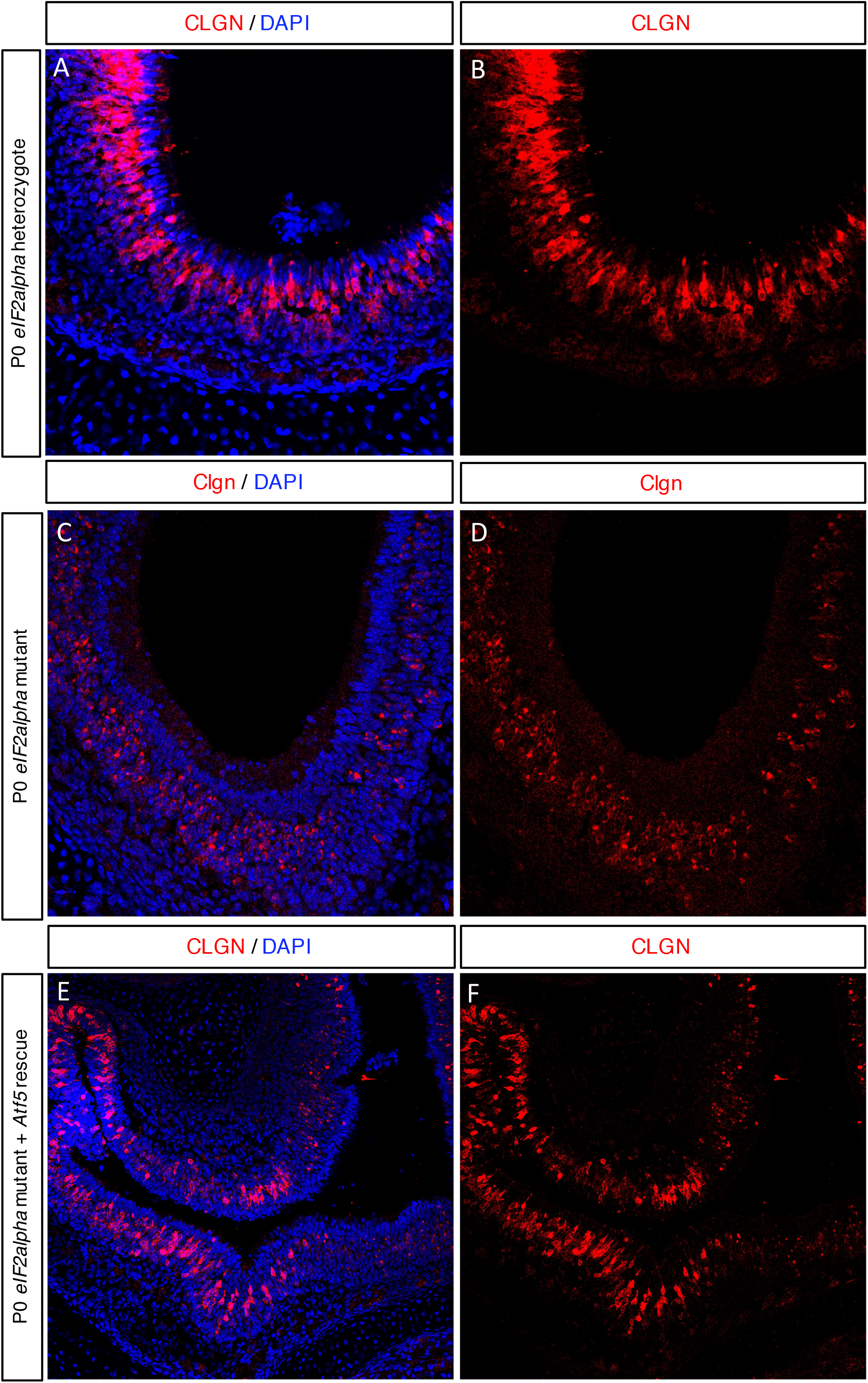
*Atf5* is required and sufficient for robust CLGN expression in the MOE. Representative LSM images. Shown are 14um coronal sections of MOE with CLGN IF in red and DAPI in blue. (A-B) A P0 *eIF2α S51A/+* control animal displaying several cell layers of robust CLGN expression. (C-D) A littermate *eIF2αS51A/S51A* animal, with reduced and more punctate CLGN expression. (E-F) An *eIF2α S51A/S51A; Gng8-tta; tetO-Atf5* transgenic rescue animal demonstrating a strong rescue of CLGN immunoreactivity in some areas and mutant levels of CLGN expression elsewhere.

### *Atf5* is Required and Sufficient for CLGN Expression in VSNs

*Atf5* was recently shown to be expressed in VSNs, where it is required for the maturation and survival of basal VSNs, and to a lesser extent apical VSNs(Nakano et al., 2016). *Atf5* translation in VSNs, as in OSNs, is driven by PERK-mediated phosphorylation of eIF2α. However, the Atf5-driven transcriptional program in VSNs has yet to be defined. In order to determine whether *Atf5* is important for CLGN expression in the VNO, I assayed CLGN immunoreactivity in VNOs harvested from the same animals used in **Figure 4**. As can be seen in **Figure 5A-B**, at P0 in eIF2α control animals CLGN is robustly expressed in the VNO, with expression levels greatest in the central (i.e. non-marginal) areas of the tissue, which correspond to mature VSNs. In contrast, eIF2α phosphomutants show a striking reduction of CLGN expression, consistent with the requirement of *Atf5* for VSN maturation. As was observed in the MOE, transgenic rescue of *Atf5* expression also rescued CLGN expression (**Figure 5E-F**). In the VNO, CLGN expression appeared to be rescued to levels greater than those observed in control animals. Thus, as was observed in the MOE, CLGN expression in the VNO is under *Atf5* control.

**Figure 5.**
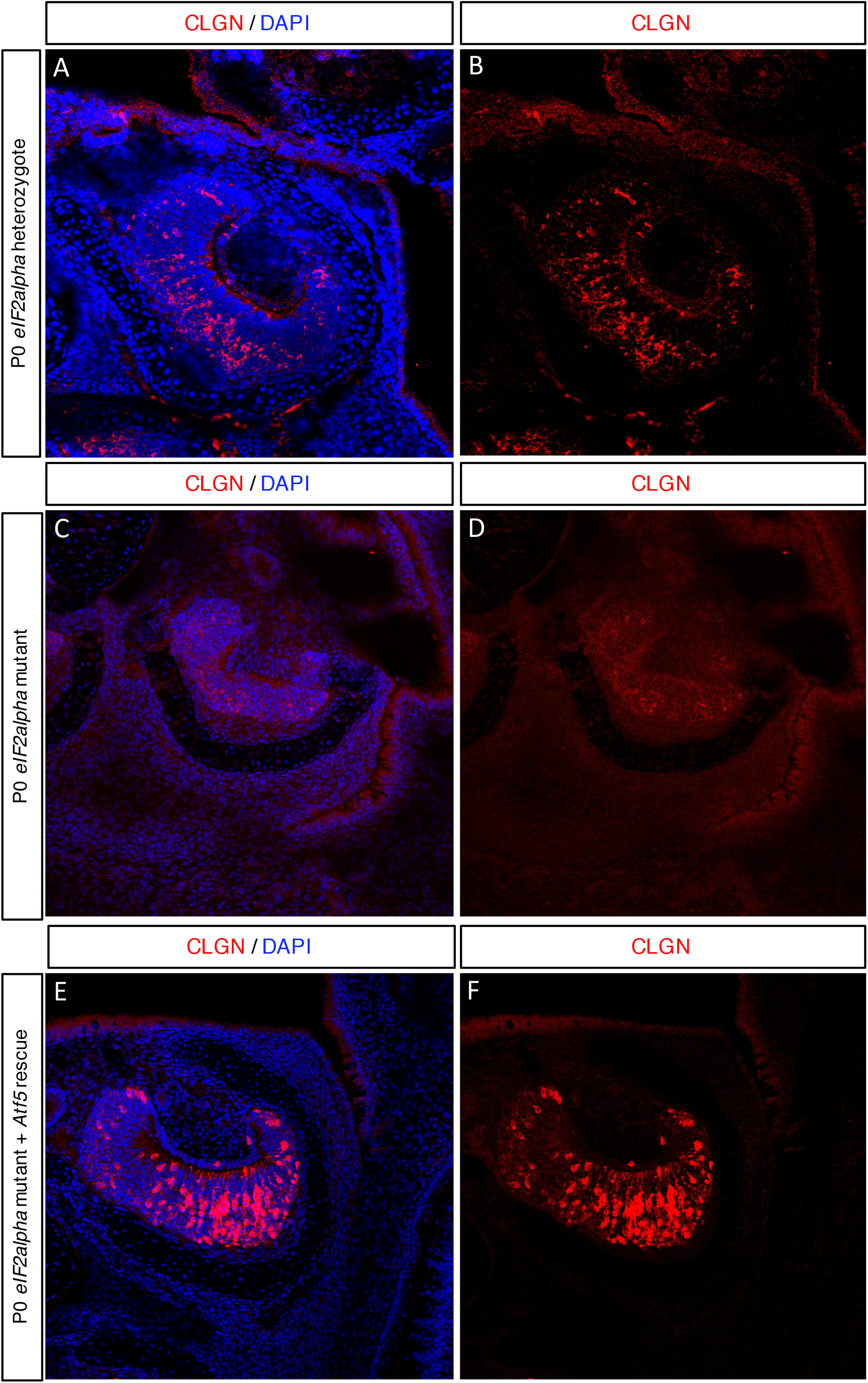
*Atf5* is required and sufficient for robust CLGN expression in the VNO. Representative LSM images. Shown are 14um coronal sections of VNO with CLGN IF in red and DAPI in blue. (A-B) A P0 *eIF2α S51A/+* control animal showing robust CLGN immunoreactivity in the central areas of the VNO. (C-D) A littermate *eIF2α S51A/S51A* animal showing loss of CLGN immunoreactivty. VNO is also decreased in size. (E-F) An *eIF2α S51A/S51A; Gng8-tta; tetO-Atf5* transgenic rescue animal showing a rescue of CLGN expression.

## Discussion

With this work, I demonstrate that Clgn, an ER chaperone required for sperm fertility, is a specific marker of mature OSNs and VSNs. These findings have a number of interesting implications. First, Clgn-deficient sperm have been shown to fail to migrate into oviducts and to be unable to bind to the zona pellucida(Yamagata et al., 2002) (Ikawa et al., 1998). These defects appear to be due to a failure of fertilin alpha/beta heterodimerization (Cho et al., 1998) (Tokuhiro et al., 2012). Fertilin alpha/beta do not appear to be involved in human spermatogenesis (Jury et al., 1997; Choi et al., 2016), limiting the translational importance of these findings. The finding that Clgn is expressed in the olfactory system raises the interesting possibility that defects in functional OR expression could also contribute to the inability of Clgn mutant sperm to migrate into oviducts. However, this possibility has yet to be directly tested.

Second, it has been widely demonstrated that chemoreceptors cannot exit the ER when expressed in cell lines, suggesting that they require a specialized biosynthetic network for their efficient production. Given that OSNs experience receptor-driven ER stress until they mature, it is likely that the full OR biosynthetic pathway is not in place until OSN maturation. Clgn is therefore an excellent candidate for membership in this pathway. Because Clgn is expressed in both OSNs and VSNs, it may aid in the folding or production of multiple classes of chemoreceptors. Alternatively, it may be involved in production of protein other than chemoreceptors. Differentiating between these possibilities is not possible without a full functional characterization of Clgn in OSNs as well as an assessment of OR expression and function in Clgn mutant animals.

Third, while receptor-transporting proteins and heat-shock proteins that increase cell-surface expression of ORs and VRs have been described, an OR-specific or VR-specific glycan-binding chaperone has not. This is of particular note for ORs, both because ORs appear to have a single, positionally-stereotyped N-terminal N-glycosylation site, and because mutation of that site prevents efficient OR feedback. Interaction with glycan-binding chaperones may therefore represent an important step in the pathway by which ORs are recognized as ORs. Whether ORs or VRs, or subsets thereof interact directly with Clgn remains to be seen. In sum, these findings identify a novel, developmentally-regulated marker of OSNs and VSNs, and suggest a compelling candidate for future studies of the specialized biosynthetic apparatuses of these cells.

